# A hydrogen-producing mitochondrion in an anaerobic eukaryotrophic rhizarian

**DOI:** 10.64898/2026.09.23.753574

**Authors:** Maggie Lawton, Yana Eglit, Courtney W. Stairs, Ryan M.R. Gawryluk

## Abstract

Diverse eukaryotes thrive under low oxygen conditions, in part through highly modified mitochondrion-related organelles (MROs) that use alternate metabolic pathways to support ATP production and cofactor recycling. Anaerobic lifestyles have evolved repeatedly across the eukaryotic tree of life, each providing an independent opportunity to understand how eukaryotes adapt to life in low oxygen conditions. Here, we use single-cell transcriptomics to reconstruct the MRO metabolism of PCE SSF, a benthic eukaryotrophic flagellate and the first cultivated representative of Novel Clade 12 (NC12; Rhizaria), an independently anaerobic rhizarian lineage. PCE SSF possesses an anaerobic hydrogen-producing mitochondrion capable of hydrogenosome-type substrate-level phosphorylation. It also retains a nearly complete but likely branched tricarboxylic acid pathway that lacks citrate synthase and malate dehydrogenase. The function of citrate synthase may instead be fulfilled by the typically cytosolic ATP citrate lyase, previously reported in this context only in the anaerobic cercozoan, *Brevimastigomonas motovehiculus*. Unlike *B. motovehiculus*, however, PCE SSF retains only Complex II and the NuoE/NuoF subunits of the electron transport chain and lacks a mitochondrial genome. Together, these features indicate an atypical and reduced mitochondrial metabolism, highlighting the diversity of evolutionary solutions to anaerobic energy metabolism in eukaryotes.

**Significance statement:** Mitochondria have repeatedly undergone extensive metabolic remodeling in eukaryotes adapted to low oxygen environments, but our understanding of this process is based on a small and phylogenetically limited sample of anaerobic lineages. Here, single-cell sequencing reveals that the recently reported protist PCE SSF possesses an unusual hydrogen-producing mitochondrion with a branched and extensively remodeled energy metabolism. Its distinctive combination of retained, repurposed, and possibly acquired metabolic pathways expands the known diversity of mitochondria and highlights how much remains to be discovered by investigating anaerobic metabolism in poorly sampled eukaryotic groups such as Rhizaria.

## Introduction

Most eukaryotes generate the bulk of their ATP via oxidative phosphorylation. In this process, electrons derived from the oxidation of organic molecules pass through the mitochondrial electron transport chain (ETC) to drive proton pumping across the inner mitochondrial membrane, establishing an electrochemical gradient that powers ATP synthesis, with oxygen acting as the terminal electron acceptor. Under low oxygen conditions, ATP can instead be generated through substrate-level phosphorylation, in which phosphate groups are directly transferred from metabolic intermediates to ADP independently of an electron transport chain. Many aerobic eukaryotes can use substrate-level phosphorylation to generate ATP and recycle redox cofactors like NADH during transient oxygen limitation, for example through ethanolic fermentation in goldfish (Van den Thillart et al. 1983). In contrast, anaerobic protists adapted for sustained growth in low-oxygen environments employ different terminal electron acceptors and rely primarily on substrate-level phosphorylation pathways distinct from those found in aerobes. These lineages have often lost components of the oxidative phosphorylation machinery and instead harbour highly modified mitochondria, termed mitochondrion-related organelles (MROs), that support anaerobic energy metabolism.

Previous literature classified mitochondria into five categories: aerobic mitochondria, anaerobic mitochondria, hydrogen-producing mitochondria, hydrogenosomes, and mitosomes (Müller et al. 2012). Anaerobic mitochondria resemble their aerobic counterparts but use alternate terminal electron acceptors such as fumarate or nitrate (Finlay et al. 1983). Hydrogenosomes, first described in *Tritrichomonas* (Lindmark and Muller 1973), have lost the tricarboxylic acid (TCA) cycle, electron transport chain (ETC), and mitochondrial genome (mtDNA), and instead produce ATP via a substrate-level phosphorylation pathway that generates molecular hydrogen as a byproduct (Carlton et al. 2007). Hydrogen-producing mitochondria possess features of both anaerobic mitochondria and hydrogenosomes, retaining a functioning electron transport chain in addition to hydrogen-generating enzymes (Müller et al. 2012). In contrast, mitosomes lack ATP-producing pathways altogether, and retain only a limited subset of mitochondrial functions, most notably iron-sulfur (Fe-S) cluster synthesis (Tovar et al. 1999; Tovar et al. 2003).

However, MRO diversity does not fit within discrete categories but instead spans a continuum of metabolic rerouting and reduction, with many lineages exhibiting combinations of traits associated with multiple MRO types (Stairs et al. 2015; Gawryluk et al. 2016). For example, *Acanthamoeba castellanii* possesses a gene-rich mitochondrial genome and complete oxidative phosphorylation (Burger et al. 1995; Gawryluk et al. 2014), yet is likely also capable of hydrogen production (Leger et al. 2013). Conversely, the MROs of *Pygsuia biforma* have hydrogenosomal enzymes but lack mtDNA and retain only a few TCA cycle and ETC enzymes such as fumarase and succinate dehydrogenase, respectively (Stairs et al. 2014).

MROs from diverse eukaryotes show similar combinations of pathway loss, rewiring of ancestral mitochondrial metabolism, and acquisition of novel functions, sometimes through lateral gene transfer (LGT) (Roger et al. 2017; Gawryluk and Stairs 2021), suggesting that adaptation to anaerobic environments has repeatedly driven convergent mitochondrial reduction across eukaryotes. As such, each independent origin of an MRO provides a valuable snapshot for understanding the evolutionary processes underlying mitochondrial adaptation to low oxygen.

Rhizaria represents a promising but comparatively understudied source of anaerobic eukaryotic diversity, as only a handful of anaerobic rhizarians have been characterized. These include the free-living cercozoan *Brevimastigomonas motovehiculus*, which combines hydrogen-producing substrate-level phosphorylation with a nearly-complete ETC and TCA cycle (Gawryluk et al. 2016), several foraminiferans that retain oxidative phosphorylation while also possessing denitrification pathways that enable anaerobic respiration with nitrate as a terminal electron acceptor (Woehle et al. 2018; Gomaa et al. 2021), one foraminiferan with an atypical anaerobic ETC (Gomaa and Bernhard 2026), and parasitic ascetosporeans that possess highly reduced mitosomes despite residing in oxygenated intracellular environments of marine invertebrates (Burki et al. 2013; Onuț-Brännström et al. 2023).

The recent cultivation of previously unknown rhizarian lineages, represented by the free-living eukaryotrophic flagellates PCE SSF and QSI PG (Eglit et al. 2024), further expands this diversity and indicates that the mitochondrial biology of anaerobic rhizarians remains substantially undersampled. PCE SSF belongs to Novel Clade 12 (NC12), a lineage otherwise known primarily from anaerobic marine and freshwater sediments (Bass et al. 2009). Here, we used single-cell sequencing to generate a transcriptome and partial nuclear genome assembly for PCE SSF and reconstruct its MRO metabolism. We identified an anaerobic mitochondrion that combines hydrogen-producing substrate-level phosphorylation with a highly reduced electron transport chain and a near-complete tricarboxylic acid pathway. The organelle also possesses mitochondrion-targeted ATP citrate lyase (ACL) and NDP-forming acetyl-CoA synthetase (ACS), enzymes that may provide additional routes for ATP generation. Together, these features suggest that the MRO of PCE SSF represents a distinct trajectory of mitochondrial reduction within Rhizaria and provides further insight into the convergent metabolic adaptations associated with anaerobic lifestyles in eukaryotes.

## Results

### Numerous mitochondrion-like organelles are present in PCE SSF

Fluorescence imaging with MitoTracker Orange revealed that PCE SSF cells contain numerous stained organelles, often collectively occupying half or more of the cell volume (Figures 1, S1). These organelles were ovaloid and numerous, with typically ∼50 per cell (37-130, n=6). Under the fixation conditions employed here, there was no visible or consistent organization of MROs within the cell relative to the disc-like nucleus or other cellular structures. As MitoTracker accumulates in response to negative membrane potential, its presence indicates that these organelles maintain a membrane potential supportive of active mitochondrial metabolism. No mitochondrial DNA was apparent with DAPI staining, but this is typical even in aerobic animal mitochondria, varying based on the quantity of mitochondrial DNA and quality of imaging (Dellinger and Gèze 2001).

**Fig. 1.**
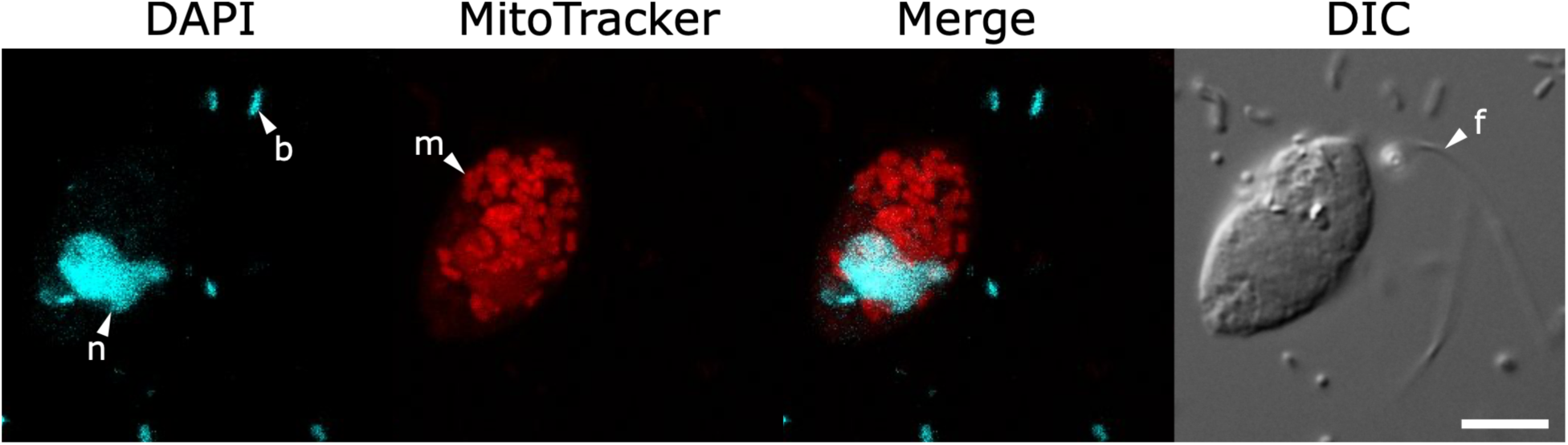
Fluorescence microscopy of PCE SSF showing DAPI (4′,6-diamidino-2-phenylindole) staining of nucleic acids and MitoTracker Orange CMTMRos staining of mitochondrion-related organelles (MROs). Images were acquired with a 100× objective. Scale bars, 5 µm. ‘n’ indicates the nuclear DNA, ‘b’ indicates bacterial DNA, ‘m’ indicates a mitochondrion-related organelle, and ‘f’ indicates a flagellum. **Alt text:** A PCE SSF cell fluorescently stained with DAPI (nucleic acids) and MitoTracker (MROs). The cell appears to have a single, irregularly-shaped nucleus, as well as many densely-packed MROs.

### Single-cell sequencing produced a reasonably complete transcriptome

The sequencing, co-assembly, and decontamination of RNA-seq reads from four individual PCE SSF cells produced a transcriptome estimated to be 56.1% complete with BUSCO (eukaryote_odb10; Table S1). This score is modest relative to those of well-characterized model eukaryotes with completely sequenced nuclear genomes, like the endomyxan *Plasmodiophora brassicae* (88.2%) (Javed et al. 2024), but BUSCO can underestimate the completeness of transcriptomes from highly divergent protists, and several missing BUSCOs in PCE SSF are associated with aerobic mitochondrial functions that are likely absent from this anaerobic lineage (Table S1). However, the BUSCO score for the PCE SSF transcriptome is comparable to that of the anaerobic rhizarian *B. motovehiculus* (54.1%) (Gawryluk et al. 2016), and significantly higher than the parasitic *M. mackini* (10.2%) (Burki et al. 2013), as listed in the EukProt database (Richter et al. 2022), supporting its suitability for reconstructing MRO metabolism. The PCE SSF single-cell genome assembly was estimated to be 16.1% complete with BUSCO and was used primarily for demonstrating eukaryotic provenance of genes based on the presence of spliceosomal introns.

The bulk transcriptome produced for PCE Brev and co-cultured bacteria was highly complete at 78.1% as estimated with BUSCO (eukaryote_odb10; Table S1), notably higher than that of other breviates such as *P. biforma* (62.4%) (Stairs et al. 2014), as reported in the EukProt database (Richter et al. 2022). Excellent coverage of the prey transcriptome allowed for thorough decontamination of the PCE SSF dataset, and high confidence that putative PCE SSF transcripts of interest did not derive from eukaryotic prey or bacteria.

### PCE SSF possesses a complex MRO without a mitochondrial genome

A total of 189 putative MRO-targeted proteins were predicted, most (167/189) of which are represented by complete transcripts. The majority of putative MRO proteins (104/189) possessed strong N-terminal MTS, while others (44/189) were membrane-associated or small proteins of known mitochondrial localization that are expected to lack MTS, such as mitochondrial carrier family proteins or protein import complex subunits. The remaining proteins (41/189) were confidently predicted to be MRO-targeted but could not be unambiguously annotated. Some transcripts (45/189) encoding predicted MRO proteins aligned to the nuclear genome assembly and contained gaps with 5ʹ GT-AG 3ʹ boundaries suggestive of spliceosomal introns. The nucleotide and amino acid sequences of predicted MRO proteins are provided in Table S2, alongside their targeting probabilities, best BLAST hits, relative abundance, and other accompanying statistics.

Despite exhaustive searches, we did not identify evidence for a mitochondrial genome in the PCE SSF metagenome assembly, or evidence of an MRO-targeted gene expression system in the transcriptome assembly, suggesting that, like many anaerobes (Carlton et al. 2007; Stairs et al. 2014; Noguchi et al. 2015), it lacks mitochondrial DNA.

### Conservation of mitochondrial transport, protein import, and Fe–S cluster assembly machineries

Many of the proteins identified in PCE SSF’s MRO are associated with conserved organellar functions, such as metabolite transport, protein import, organellar maintenance, and Fe-S cluster synthesis. Among these were numerous transport proteins mediating metabolite exchange across the MRO membrane. We identified 19 members of the mitochondrial carrier protein family, including an ADP/ATP translocase, a phosphate transporter, and multiple other carriers whose substrate specificities could not be confidently predicted (Figure 2).

**Fig. 2.**
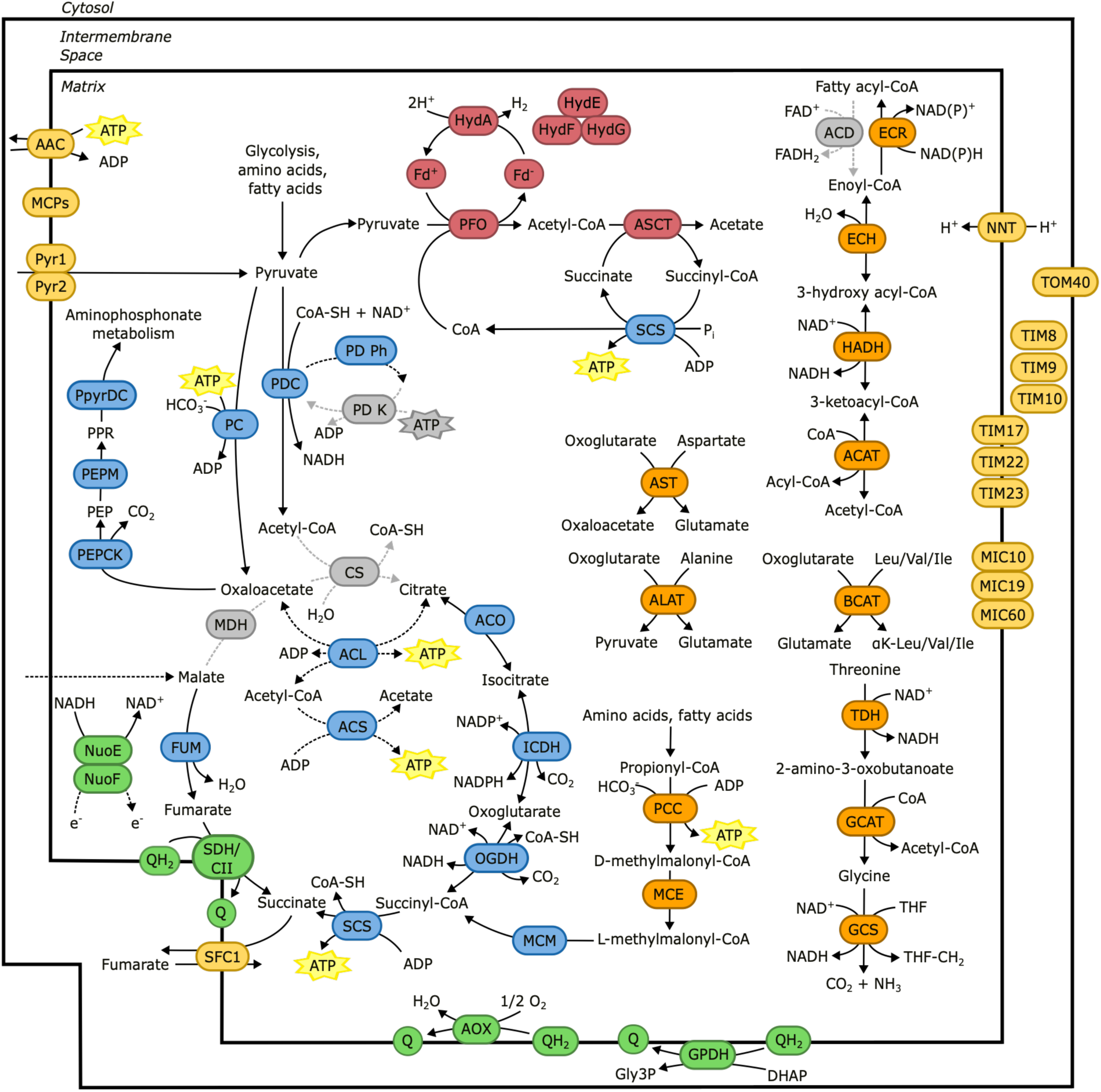
Reconstruction of PCE SSF MRO metabolism, highlighting pathways associated with ATP production. Arrows indicate expected directions of reactions: solid arrows indicate well-described reactions, while dashed arrows indicate reactions theorized in this paper, and grey dashed arrows indicate absent reactions. The question mark connected to NuoE/NuoF indicates that the oxidation mechanism is undetermined. Bubbles indicate predicted MRO-targeted proteins, and color indicates associated pathway: Red (hydrogen-producing substrate-level phosphorylation), pyruvate metabolism (blue), amino acid and fatty acid metabolism (orange), membrane transport (yellow), electron-transport (green), absent (grey). Abbreviations: 2-amino-3-ketobutyrate CoA ligase (GCAT), 3-hydroxyacyl-CoA dehydrogenase (HADH), acetate:succinate Co-A transferase (ASCT), acetyl CoA acetyltransferase (ACAT), acetyl-CoA synthetase (ACS), aconitase (ACO), acyl-CoA dehydrogenase (ACD), ADP/ATP translocase (AAC), alanine aminotransferase (ALAT), alternative oxidase (AOX), aspartate aminotransferase (AST), ATP citrate lyase (ACL), branched-chain amino acid aminotransferase (BCAT), citrate synthase (CS), Complex I subunit NuoE (NuoE), Complex I subunit NuoF (NuoF), enoyl-CoA hydratase (ECH), ferredoxin (Fd^+/-^), fumarase (FUM), glycerol-3-phosphate dehydrogenase (G3PDH), glycine cleavage system (GCS), [FeFe]-hydrogenase (HydA), HydA maturases (HydE, F, G), malate dehydrogenase (MDH), methylmalonyl-CoA epimerase (MCE), methylmalonyl-CoA mutase (MCM), MICOS complex subunit (MIC), mitochondrial carrier protein (MCP), NAD(P) transhydrogenase (NNT), oxoglutarate dehydrogenase (OGDH), pyruvate carboxylase (PC), pyruvate dehydrogenase complex (PDC), PDC kinase (PDC K), PDC phosphatase (PD Ph), propionyl Co-A carboxylase (PCC), phosphoenolpyruvate (PEP), PEP carboxykinase (PEPCK), PEP mutase (PEPM), 3-phosphonopyruvate (PPR), phosphonopyruvate decarboxylase (PpyrDC), pyruvate carrier 1/2 (Pyr1/2), pyruvate:ferredoxin oxidoreductase (PFO), quinol (QH2), quinone (Q), succinate dehydrogenase/Complex II (SDH/CII), succinyl-CoA synthetase (SCS), threonine dehydrogenase (TDH), trans-2-enoyl-CoA reductase (ECR), translocase of the inner membrane (TIM), translocase of the outer membrane (TOM). **Alt text:** Metabolic diagram outlining the key pathways associated with ATP production in the MRO of PCE SSF. The MRO appears to be depleted of some oxygen-associated pathways, including most of the ETC and CS and MDH of the TCA cycle.

PCE SSF MROs possess most core components of the mitochondrial protein import machinery, consistent with the robust *in silico* prediction of mitochondrial targeting sequences. Identified components of the protein import apparatus include the TOM complex (Tom7, Tom22, and Tom40), small TIM chaperones (Tim8, Tim9, and two copies of Tim10), TIM22 (Tim22), and the TIM23 complex (Tim17, Tim23, Tim14, Pam16, and Hsp70) (Figure 2). The mitochondrial processing peptidase (MPP⍺, β) was also identified, indicating that imported proteins undergo cleavage of their N-terminal targeting sequences, as in other mitochondria. No components of the SAM complex, responsible for the insertion of β-barrel proteins into the outer mitochondrial membrane (Neupert and Herrmann 2007), were identified, despite the presence of its substrates like Tom40 and a putative porin.

Divergent homologs of the core MICOS subunits Mic10, Mic19, and Mic60 were identified (Figure 2). The MICOS complex is best known for organizing crista junctions and maintaining mitochondrial architecture, but has been lost in many, although not all, acristate MROs (Muñoz-Gómez et al. 2015; Yi et al. 2026). Because the ultrastructure of the PCE SSF MRO has not yet been resolved by transmission electron microscopy, it remains unknown whether these organelles retain cristae or whether MICOS fulfils alternative roles.

PCE SSF MROs retain the canonical mitochondrial ISC (iron-sulfur cluster) pathway responsible for Fe-S cluster biosynthesis, one of the most highly conserved functions of mitochondria and MROs. Core ISC components identified include the scaffold protein IscU, the assembly factor Isca1, the cochaperone HscB, and the cysteine desulfurase Nfs1. We also identified the conserved transporter ABCB7/ATM1, which exports an unknown sulfur-containing intermediate required for cytosolic Fe-S cluster assembly (CIA) (Kispal et al. 1999; Li et al. 2022). The retention of this pathway is consistent with the numerous Fe-S cluster-containing proteins predicted to localize to PCE SSF MROs, including pyruvate:ferredoxin oxidoreductase (PFO), aconitase (ACO), and components of respiratory Complexes I and II (see below).

### PCE SSF MROs retain diverse biosynthetic and catabolic pathways

Numerous enzymes involved in amino acid metabolism were predicted to localize to MROs, including enzymes involved in the degradation of alanine, aspartate, threonine, glycine, leucine, valine, and isoleucine (Figure 2). The products of these pathways, including acetyl-CoA, oxaloacetate, and propionyl-CoA, are predicted to feed into downstream energy metabolism in PCE SSF. In contrast, MRO-associated amino acid metabolism in some anaerobic protists is more restricted; for example, the *Mastigamoeba balamuthi* hydrogenosome appears to retain primarily enzymes for glycine/serine interconversion and associated one-carbon metabolism (Gill et al. 2007; Nývltová et al. 2015).

The organelles also possess a capacity for lipid metabolism, including monoacylglycerol fatty acid metabolism, cardiolipin synthesis, and most of Type II fatty acid synthesis and ß-oxidation (Figure 2). Notably, acyl-CoA dehydrogenase (ACD), which catalyzes the initial conversion of fatty acyl-CoA into enoyl-CoA during ß-oxidation, was not detected, although trans-2-enoyl CoA reductase (ECR), which catalyzes the reverse reaction, was. The absence of ACD raises the possibility that ß-oxidation has been lost or employs an alternative enzyme, potentially favouring fatty acid synthesis over degradation. Curiously, ACD is also absent from *P. biforma* despite the presence of several other ß-oxidation enzymes (Stairs et al. 2014). If ß-oxidation is indeed active, the resulting acetyl-CoA can feed into other energy-producing pathways, discussed further below.

No complete pathway for nucleotide biosynthesis was identified in the genome-less MRO of PCE SSF. This contrasts with aerobic mitochondria, which play important roles in both cytosolic and organellar nucleotide metabolism (MacVicar 2025), but is consistent with the reduced demand for nucleotides in an organelle lacking mitochondrial DNA. Despite this, two MRO-targeted copies of dihydropyrimidine dehydrogenase (DPYD) suggest some capacity for uracil catabolism. More unexpectedly, PCE SSF encodes both cytosolic and MRO-targeted copies of ribose-phosphate pyrophosphokinase (RPPK), which converts ribose 5-phosphate into phosphoribosyl pyrophosphate (PRPP), a precursor for both purine synthesis and salvage. RPPK is typically cytosolic, and mitochondrially targeted versions have only been found in spinach (Krath & Hove-Jensen, 1999) and *Blastocystis* (Gentekaki et al. 2017). However, unlike PCE SSF, both taxa retain mitochondrial genomes together with a mitochondrial purine synthesis pathway in which RPPK can participate. As such, the function of an MRO-targeted RPPK in PCE SSF remains unclear.

### Pyruvate metabolism supports hydrogen and ATP production

Pyruvate is likely imported into PCE SSF MROs from the cytosol via the conserved pyruvate importer complex MPC1/2 (Pyr1/2) (Bricker et al. 2012), although it may also be synthesized within the organelle by alanine aminotransferase. Within the MRO, it can be converted to oxaloacetate by pyruvate carboxylase (PC), or to acetyl-CoA by either the pyruvate dehydrogenase complex (PDC) or pyruvate:ferredoxin oxidoreductase (PFO) (Figure 2). During the latter reaction, PFO transfers electrons to ferredoxin, which is subsequently reoxidized by [FeFe]-hydrogenase to generate molecular hydrogen, a hallmark of hydrogen-producing mitochondria and hydrogenosomes. Although both enzymes catalyze the conversion of pyruvate to acetyl-CoA, the coexistence of PDC and PFO has been reported in MROs from several other protists (Stechmann et al. 2008; Gawryluk et al. 2014; Gawryluk et al. 2016). The markedly higher expression of PFO relative to PDC subunits in our dataset suggests that PFO may represent the major route of pyruvate catabolism, providing both acetyl-CoA and reduced ferredoxin for downstream ATP and hydrogen production. Although multiple PFO and [FeFe]-hydrogenase homologues are encoded in the PCE SSF transcriptome, only a subset is predicted to localize to the MRO (Figures S2, S3). Phylogenetic analyses indicate that these paralogs have complex evolutionary histories, consistent with previous reports for PFO and [FeFe]-hydrogenase in anaerobic eukaryotes (Figures S2, S3).

ATP can be generated from acetyl-CoA by at least two pathways in PCE SSF MROs. In the first, acetyl-CoA is converted to acetate and succinyl-CoA by a type B1 acetate:succinate CoA-transferase (ASCT). Succinyl-CoA is then converted back to succinate by succinyl-CoA synthetase (SCS), generating ATP by substrate-level phosphorylation. Together with PFO and [FeFe]-hydrogenase, the ASCT/SCS pathway constitutes the canonical hydrogen-producing ATP-generating pathway found in many anaerobic eukaryotes. As in other anaerobic eukaryotes, these enzymes have complex evolutionary histories and are ultimately of bacterial origin, although their acquisition by eukaryotes like PCE SSF likely involved multiple eukaryote-to-eukaryote transfer events (Hug et al. 2010; Roger et al. 2017).

PCE SSF also encodes an MRO-targeted NDP-forming acetyl-CoA synthetase (ACS), providing a second route for ATP production through the conversion of acetyl-CoA to acetate. Unlike the more widespread AMP-forming ACSs, which consume ATP during acetyl-CoA synthesis, NDP-forming ACSs can operate in the opposite direction to generate ATP through substrate-level phosphorylation, as recently confirmed in anaerobic fornicates (Koganemaru et al. 2026). Similar MRO-targeted NDP-forming ACS homologs have been identified in several other anaerobic protists, including *Cantina marsupialis* (Noguchi et al. 2015) (Figure 3). Phylogenetic analysis further suggests that this class of eukaryotic ACS is most closely related to homologs from Heimdallarchaeota (Figure S4), mirroring the recent proposal that ATP-generating ACS in anaerobic fornicates was acquired by LGT from Lokiarchaeota (Koganemaru et al. 2026), although the evolutionary events underlying its distribution among anaerobic eukaryotes are uncertain.

**Fig. 3.**
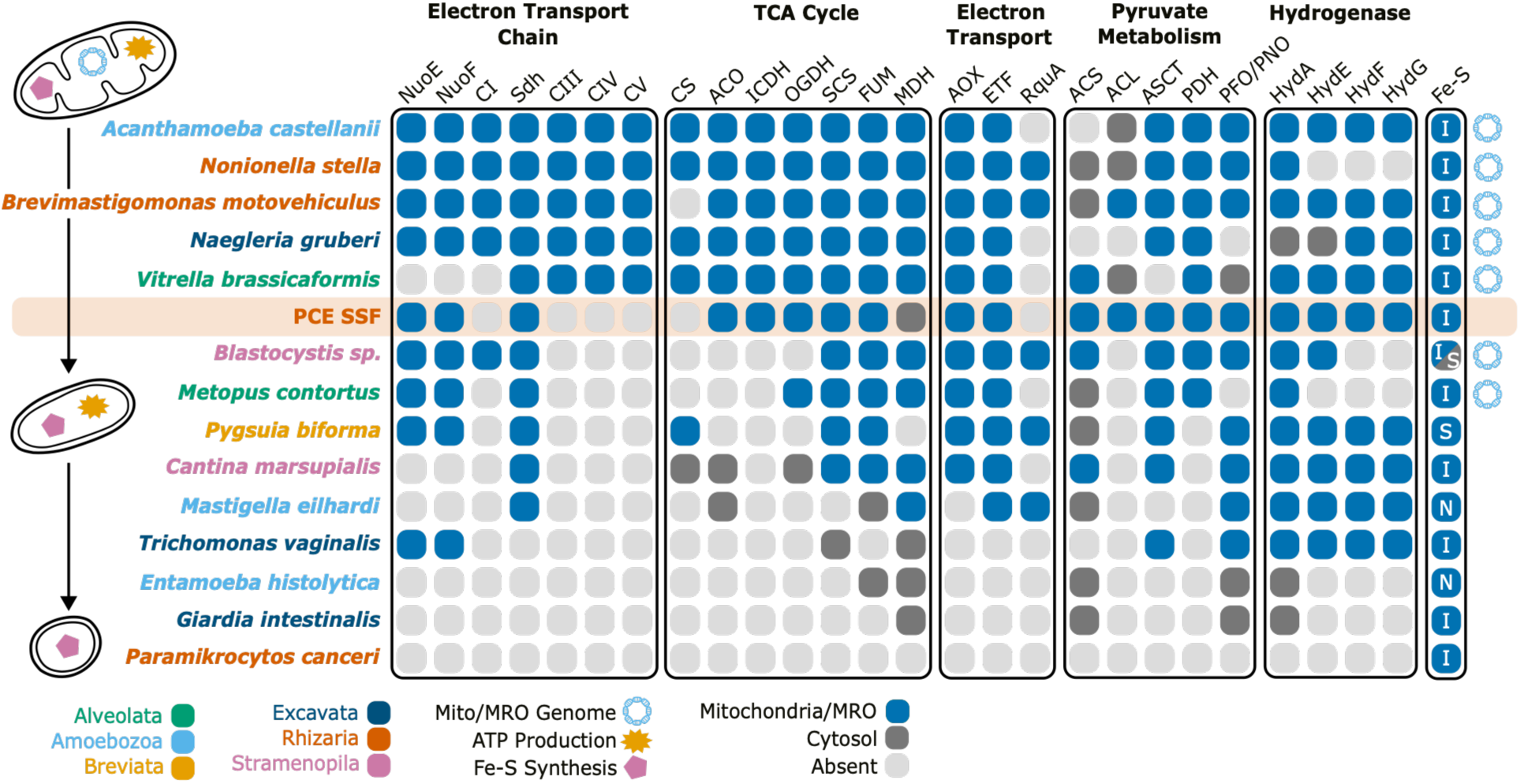
Distribution of key MRO-associated proteins across representative anaerobic protists from diverse eukaryotic lineages. Taxa are ordered vertically from least to most reduced, with icons on the left depicting a ‘typical’ aerobic mitochondrion, hydrogenosome, and mitosome, respectively. The Fe-S column indicates type of Fe-S cluster synthesis pathway present: ISC (I), SUF-like minimal system (S), or NIF (N). Abbreviations: Complex I subunit NuoE (NuoE), Complex I subunit NuoF (NuoF), Complex I (CI), succinate dehydrogenase/Complex II (SDH), Complex III (CIII), Complex IV (CIV), Complex V (CV), citrate synthase (CS), aconitase (ACO), isocitrate dehydrogenase (ICDH), oxoglutarate dehydrogenase (OGDH), succinyl-CoA synthetase (SCS), fumarase (FUM), malate dehydrogenase (MDH), alternative oxidase (AOX), electron-transferring flavoprotein (ETF), rhodoquinone synthesis protein (RquA), acetyl-CoA synthetase (ACS), ATP-citrate lyase (ACL), acetate:succinate CoA transferase (ASCT), pyruvate dehydrogenase (PDH), pyruvate:ferredoxin oxidoreductase (PFO), pyruvate:NADP^+^ oxidoreductase (PNO), [FeFe]-hydrogenase (HydA), HydA maturases (HydE, F, G). **Alt text**: Depiction of the presence, absence, and localization of key MRO-associated proteins across 15 taxonomically diverse anaerobic protists, including PCE SSF. Taxa range from highly complete to highly reduced, with each demonstrating a unique composition of proteins. Degree of completeness does not appear to follow taxonomic origin.

### An incomplete and branched tricarboxylic acid pathway

In addition to its role in anaerobic ATP production, acetyl-CoA may also enter the TCA cycle, which in aerobic mitochondria serves as a major source of NADH and other reducing equivalents for the electron transport chain and as a hub for production of biosynthetic intermediates. Although anaerobic MROs often retain only a small subset of TCA cycle enzymes, PCE SSF possesses all canonical enzymes except for citrate synthase (CS) and malate dehydrogenase (MDH) (Figure 3). This contrasts with most other MROs, in which enzymes of the oxidative branch (from citrate to oxoglutarate) are more frequently lost (Denoeud et al. 2011; Stairs et al. 2014; Noguchi et al. 2015; Stairs et al. 2015; Gawryluk et al. 2016), raising the question of how citrate is synthesized and how carbon flows through the pathway in the absence of CS and MDH.

One possible explanation is that citrate is synthesized through an alternative route. However, no additional CS homolog was identified in the transcriptome, arguing against import of citrate generated by an alternatively targeted CS. Instead, we identified a highly expressed, MRO-targeted ATP citrate lyase (ACL), raising the possibility that it compensates for the absence of CS. ACL normally catalyzes the ATP-dependent cleavage of citrate to acetyl-CoA and oxaloacetate during cytosolic fatty acid biosynthesis. In principle, however, ACL could operate in reverse to synthesize citrate while producing ATP, although it is unclear if this would be energetically favourable. The presence of ACL is notable because the only other described organism with mitochondrion-localized ACL is another rhizarian, *B. motovehiculus*, which also possesses hydrogen-producing mitochondria with a nearly complete TCA cycle lacking CS under anaerobic growth conditions (Gawryluk et al. 2016). We suspect that mitochondrial targeting evolved independently in these lineages, consistent with phylogenetic analyses of ACL (Figure S5), in which *B. motovehiculus* groups confidently within Rhizaria, while PCE SSF branches just outside of Rhizaria, sister to cryptomonad *Goniomonas*.

An alternative possibility is that ACL retains its canonical citrate-cleaving activity, while the oxidative branch of the TCA cycle operates in reverse. In this scenario, oxoglutarate generated by oxoglutarate dehydrogenase (OGDH) would be converted to isocitrate by NADP-dependent isocitrate dehydrogenase (ICDH), oxidizing NADPH, before aconitase (ACO) converts isocitrate to citrate. ACL could then cleave citrate to generate acetyl-CoA and oxaloacetate. The absence of MRO-targeted MDH, however, complicates the model. Two MDH homologs were identified in PCE SSF, but they are instead predicted to localize to peroxisomes, and therefore OAA cannot directly enter the reversed reductive branch of the TCA cycle. Instead, it may be converted to phosphoenolpyruvate by MRO-targeted phosphoenolpyruvate carboxykinase (PEPCK), thereby diverting carbon into aminophosphonate metabolism through the sequential action of phosphoenolpyruvate mutase and phosphonopyruvate decarboxylase. The same enzymes are present in the hydrogen-producing mitochondria of *B. motovehiculus* (Gawryluk et al. 2016). Ultimately, we cannot distinguish between these two models, but in either case, carbon flow through the TCA pathway is likely branched in PCE SSF MROs, as it is in some anaerobic bacteria (Amador-Noguez et al. 2010).

### A partial electron transport chain supports redox balance

PCE SSF retains a complete Complex II (CII), which is predicted to act as a fumarate reductase, reducing fumarate to succinate while reoxidizing quinols (Gawryluk and Stairs 2021). Together with fumarase (FUM), which likely acts on malate imported into the MRO, this pathway contributes to redox cofactor recycling and succinate production. The succinate produced can then act as a CoA acceptor for ASCT during ATP production (see above). In anaerobic mitochondria and MROs, fumarate reduction by CII is typically coupled to rhodoquinone (RQ), whose lower redox potential makes it well suited for electron transfer to fumarate (Gawryluk and Stairs 2021). However, despite extensive searches, only a few components of the ubiquinone synthesis pathway (COQ1, COQ8, and COQ9) were identified, none with evidence of mitochondrial targeting. In addition, no recognizable rhodoquinone biosynthesis enzyme was detected, such as RquA, which converts UQ to RQ. Thus, although PCE SSF retains some quinone biosynthetic machinery, the source of quinones used by the MRO remains unclear. One possibility is that quinones are acquired from prey; notably, RquA is present in the prey transcriptome. Similar quinone scavenging has been proposed in other heterotrophic anaerobes (Stairs et al. 2014; Stairs et al. 2018).

Unlike CII, CI is represented only by the soluble NuoE and NuoF subunits. These proteins can accept electrons from NADH, but the membrane-associated subunits required for quinone reduction by CI are absent. Similar partial CI complexes occur in other anaerobic protists (Hrdy et al. 2004; Stairs et al. 2014; Stairs et al. 2021) (Figure 3), suggesting that these proteins perform functions independent of oxidative phosphorylation. One proposed role is participation in a soluble, trimeric, electron-bifurcating complex with [FeFe]-hydrogenase, coupling NADH oxidation to hydrogen production (Boxma et al. 2007; Schut and Adams 2009; Gawryluk and Stairs 2021). Although CI is unlikely to contribute directly to quinone reduction, the MRO retains several additional enzymes predicted to exchange electrons with a quinone pool, including CII, glycerol-3-phosphate dehydrogenase (GPDH), and alternative oxidase (AOX). Electron-transferring flavoprotein (ETF) was identified, but electron-transferring flavoprotein:ubiquinone oxidoreductase (ETF-QO), which typically transfers electrons from reduced ETF to the quinone pool, was not detected. The pathway for reoxidation of reduced ETF in PCE SSF therefore remains unclear. The retention of AOX may permit oxidation of reduced quinones during transient oxygen exposure, as proposed for other anaerobic eukaryotes (Tsaousis et al. 2018). Together, these observations argue that the remaining components of the ETC primarily support redox balance rather than canonical oxidative phosphorylation.

## Discussion

### Adaptation to anoxia follows common patterns of reduction, repurposing, and lateral gene transfer

Eukaryotes have independently adapted to low oxygen environments many times, often converging on similar metabolic repertoires despite their distant evolutionary relationships. Adaptation to anaerobiosis often follows a broadly predictable trajectory in which oxygen-dependent pathways are progressively lost while oxygen-independent metabolic pathways are acquired. The extent and nature of mitochondrial reduction appear to reflect adaptation to local ecological conditions more strongly than phylogenetic relatedness alone (Figure 3). For example, highly reduced parasitic lineages such as *Giardia intestinalis*, *Entamoeba histolytica*, and *Paramikrocytos canceri* possess mitosomes incapable of ATP production, whereas free-living protists inhabiting oxic or intermittently hypoxic environments like *A. castellanii*, *Nonionella stella*, and *Naegleria gruberi* retain complete oxidative phosphorylation pathways while also encoding enzymes typically associated with hydrogenosomes. Rhizarian MROs span this continuum, and PCE SSF expands it further by representing a distinct metabolic state in which the oxygen-dependent components of the electron transport chain have been lost, while many of the TCA pathway enzymes are retained, albeit with likely altered metabolic roles.

Anaerobic eukaryotes employ a diverse array of substrate-level phosphorylation pathways, including wax ester fermentation in euglenids (Inui et al. 1982) and arginine dihydrolysis in *G. intestinalis* (Schofield et al. 1990). Like many other anaerobic protists, PCE SSF relies predominantly on hydrogenosome-type ATP production (Figure 3) but likely supplements its ATP production through additional pathways involving ACS and potentially ACL. The coexistence of these partially overlapping pathways highlights the metabolic flexibility that has repeatedly evolved in anaerobic MROs. Although the evolutionary origins of the hydrogenosomal enzymes are still debated (Martin and Müller 1998; Degli Esposti et al. 2016), accumulating evidence favours repeated lateral acquisition from prokaryotes followed by extensive transfer between anaerobic eukaryotes, rather than strict inheritance from a facultatively anaerobic mitochondrial ancestor (Hug et al. 2010; Roger et al. 2017). Consistent with this, the MRO-targeted PFO and [FeFe]-hydrogenase of PCE SSF are most similar to eukaryotic homologs, but do not specifically group with homologs from other anaerobic rhizarians, indicating that they were independently recruited into its anaerobic metabolism (Figures S2, S3).

### PCE SSF possesses a distinctive anaerobic hydrogen-producing mitochondrion

The metabolic reconstruction detailed here reveals an anaerobic hydrogen-producing mitochondrion in PCE SSF that combines canonical hydrogenosomal ATP production with the retention of large portions of mitochondrial carbon metabolism. Among described anaerobic eukaryotes, PCE SSF occupies what appears to be a unique metabolic position, as no others retain the oxidative branch of the TCA pathway (ACO, ICDH, OGDH) while lacking respiratory Complexes III-V (Figure 3). Rather than representing a simple reduction of ancestral mitochondrial metabolism, the retained pathways appear to have been reorganized. ACL may either compensate for the absence of citrate synthase or, alternatively, support reverse flux through the oxidative branch of the TCA pathway.

Its capacity for predation also places PCE SSF in a unique metabolic and ecological niche. The absence of a recognizable complete quinone synthesis pathway, despite the retention of several MRO-localized quinone-dependent enzymes, raises the possibility that quinones are scavenged from prey, as has been proposed for other anaerobic eukaryotes (Stairs et al. 2018; Salomaki et al. 2021). PCE SSF is also, to our knowledge, one of the first described anaerobic eukaryotrophs. It was previously suggested that anaerobic eukaryotrophy might be energetically prohibitive because substrate-level phosphorylation alone yields relatively few ATP molecules (Fenchel 2011). However, the discovery of PCE SSF and QSI PG (Eglit et al. 2024) demonstrates that eukaryotrophy can be successfully coupled to highly modified anaerobic mitochondria. In PCE SSF, this has been achieved through extensive metabolic reorganization in addition to progressive pathway loss.

## Methods

### Cultivation of PCE SSF

PCE SSF was maintained at room temperature (18-25°C) in 4% (v/v) LB in sterile seawater in co-culture with its prey, the undescribed breviate PCE Brev. PCE Brev was maintained on a polyxenic assemblage of co-cultured bacteria. PCE SSF cultures were maintained through weekly transfers into 15 mL conical tubes containing 12 mL of medium pre-inoculated one week earlier with PCE Brev and its associated bacterial community.

### PCE SSF cell picking and sequencing

Despite existing as a stable culture, it was not possible to grow PCE SSF to a sufficient density for bulk sequencing. Instead, single cells were manually isolated under a Nikon Ti-2A inverted microscope using a finely drawn Pasteur pipette, washed three times in spent medium sterilized by passage through a 0.22 µm cellulose acetate filter, and either transferred into 2 µL of water (for genome amplification) or 0.2 µL SUPERase·In RNase Inhibitor (20 U/μL; ThermoFisher Scientific) and 1.8 µL 0.2% (v/v) Triton X-100 (for transcriptome amplification). Cells were stored at −80°C and lysed via three freeze-thaw cycles in liquid nitrogen.

For single-cell transcriptomics, the Smart-seq2 protocol (Picelli et al. 2014) was used to generate cDNA from picked cells. Sequencing libraries were then prepared from cDNA derived from four individual cells at the UBC Sequencing and Bioinformatics Consortium (Vancouver, BC, Canada) and 2 x 150 bp PE reads were sequenced on an Illumina NextSeq 500 platform. Sequencing statistics are provided in Table S1.

To generate a single-cell metagenome assembly, DNA from picked PCE SSF cells was amplified using the 4BB TruePrime Single Cell WGA Kit (4basebio) following the manufacturer’s protocol with the following modification: the final incubation at 30°C was increased to 6 h. PCR-free shotgun DNA sequencing libraries were generated at Génome Québec from four individual cells and 2 x 150 bp reads were sequenced on a NovaSeq PE150 platform (Illumina). Sequencing statistics are provided in Table S1.

### PCE Brev transcriptome and metagenome sequencing

To eliminate the possibility that candidate PCE SSF MRO proteins are derived from co-cultured microbes, we generated a high-quality bulk transcriptome from PCE Brev grown without PCE SSF for use in bioinformatic decontamination. PCE Brev was grown in 4% LB in seawater in unvented 75 cm^2^ tissue culture flasks pre-inoculated with *E. coli* to deplete oxygen from the medium. After one week, a 12 mL volume of PCE Brev culture was added to each flask and cells were allowed to grow for another week. For RNA extraction, each flask was emptied and rinsed in autoclaved seawater to remove excess bacteria. The bottom surface, retaining PCE Brev and adherent bacteria, was scraped in 10 mL of autoclaved seawater and transferred into a 15 mL conical tube. Cells were centrifuged at 1,500 x *g* for 10 min and the supernatant was discarded. Pellets were resuspended in seawater, combined, and centrifuged again at 1,500 x *g* for 5 min. A ∼200 mg pellet was combined with 800 µL of TRIzol reagent (ThermoFisher Scientific) and RNA was extracted following the manufacturer’s protocol. RNA was precipitated in a 1 M lithium chloride solution overnight at 4°C. A NEBNext Ultra II Directional poly-A+ mRNA Library Prep sequencing library was generated at The Centre for Applied Genomics (The Hospital for Sick Children, Toronto, ON, Canada) and 2 x 150 bp PE reads were sequenced on a NovaSeq X platform (Illumina).

### Sequence Assembly and Quality Control

RNA-seq reads from all four cells of PCE SSF were visually inspected for quality using FastQC (v0.12.0; Andrews 2010), combined, trimmed with Trimmomatic (v0.30; Bolger et al. 2014) and assembled using Trinity (v.2.14.0; Grabherr et al., 2011), with default settings and custom trimming references for the Smart-seq2 primers. Bulk transcriptome reads from PCE Brev were similarly trimmed and assembled using Trinity, with default settings.

Assembled PCE SSF transcripts were used to query the PCE Brev transcriptome with blastn (v2.13.0; Altschul et al., 1990) to identify sequences derived from PCE Brev and co-cultured bacteria. All sequences with a match of ≥95% nucleotide identity over ≥100 bp length were removed with SeqKit grep (v0.15.0; Shen et al., 2016). The PCE SSF transcriptome was then compared to the NCBI nr database (downloaded Nov 7, 2022) using DIAMOND blastx (v2.0.15; Buchfink et al., 2021). All sequences with bacterial matches ≥ 90% or matches of ≥ 95% identity to *Talaromyces*, a minor fungal contaminant identified in the dataset, were discarded if the alignment was ≥ 100 amino acids in length. The assembled and decontaminated PCE SSF transcriptome was then trimmed to remove remnants of the Smart-seq2 locked nucleic acid primer sequence using CutAdapt (v4.6; Martin 2011), and open reading frames were predicted and translated using TIdeS (v1.3.5; Maurer-Alcalá & Kim 2024).

For assembly of a single-cell metagenome, PCE SSF genomic reads were normalized with BBNorm (v38.86; https://sourceforge.net/projects/bbmap/) to reduce assembly time and complexity and trimmed with fastp (v0.23.4; Chen et al. 2018). Reads from all four cells were combined and assembled using metaSPAdes (v3.15.4; Nurk et al. 2017; Prjibelski et al. 2020). Genome quality statistics were estimated with QUAST (v5.0.2; Gurevich et al., 2013). The metagenome assembly was queried against the PCE Brev metagenome (Aguilera-Campos et al. 2025) with blastn (v2.13.0; Altschul et al. 1990), and all sequences with a match of ≥95% nucleotide identity over ≥100 bp length were removed with SeqKit grep (v0.15.0; Shen et al. 2016). Complete assembly statistics are listed in Table S1.

Completeness of the genome and transcriptome assemblies was estimated with BUSCO using the eukaryote_odb10 dataset (v5.5.0; Manni et al. 2021). Transcript relative abundance was estimated using Salmon (v1.7.0; Patro et al. 2017) and bowtie2 (v2.4.1; Langmead & Salzberg 2012) through Trinity’s align_and_estimate_abundance.pl script.

To assess the presence of a mitochondrial genome, 16S rRNA gene sequences were predicted from the PCE SSF metagenome assembly using Barrnap (v0.9; https://github.com/tseemann/barrnap) and inspected for evidence of mitochondrial genome provenance. In addition, we searched the metagenome for components of the mitochondrial gene expression machinery, including mitochondrial ribosomal proteins.

### MRO Proteome Prediction and Analysis

MRO proteins were predicted based on homology to sequences in previously published mitochondrial/MRO datasets and identification of mitochondrial targeting sequences (MTS) in predicted protein N-termini. Mitochondrial targeting and localization were predicted using the alpha release of CoMR (Boisard et al. 2026), utilizing TargetP (v2.0; Armenteros et al. 2019), MitoProt II (v1.101; Claros & Vincens 1996), MitoFates (metazoa and fungi; Fukasawa et al. 2015), and DeepLoc (v2.0; Thumuluri et al. 2022). Sequences exceeding the selected targeting probability thresholds (>0.5 MitoFates, >0.75 TargetP, >0.9 MitoProt II) were retained. These thresholds followed published recommendations for MitoFates, whereas more stringent thresholds were applied to TargetP and MitoProt II based on manual curation. DeepLoc predictions were not used for classification because predictions for short protein sequences were frequently inconsistent with other evidence. Sub-mitochondrial localization was predicted, where possible, using DeepMito (Savojardo et al. 2020). Predicted MRO protein sequences lacking identifiable MTS, either because they were incomplete or not expected to have MTS, were identified through homology to the curated MRO proteomes of *B. motovehiculus* (Gawryluk et al. 2016)*, A. castellanii* (Leger et al. 2013)*, P. biforma* (Stairs et al. 2014), and *Blastocystis hominis* (Stechmann et al. 2008). All sequences were then manually curated to confirm their identity and predicted targeting, and to remove residual contaminants.

To align transcripts encoding predicted MRO proteins to the metagenome assembly, we used Exonerate (v2.4.0; Slater & Birney 2005) with the cdna2genome alignment model and retaining only alignments with at least 40% of the maximal score for that query. Alignments demonstrating the presence of sequence gaps in transcripts relative to their genomic counterpart, separated by 5ʹ GT-AG 3ʹ boundaries, were taken as evidence of canonical spliceosomal introns and the eukaryotic origin of that gene sequence.

For comparative analyses, the presence and subcellular localization of key MRO-associated metabolic proteins were assessed across representative anaerobic protists. Homologs were identified in publicly available sequence databases using blastp (v2.13.0; Altschul et al. 1990), and mitochondrial targeting was assessed as described above. Protein presence and localization were further evaluated against previously published genomic, transcriptomic, and localization data (Anderson and Loftus 2005; Carlton et al. 2007; Fritz-Laylin et al. 2010; Emelyanov and Goldberg 2011; Leger et al. 2013; Gawryluk et al. 2014; Stairs et al. 2014; Tsaousis et al. 2014; Flegontov et al. 2015; Noguchi et al. 2015; Stairs et al. 2015; Gawryluk et al. 2016; Gentekaki et al. 2017; Lewis et al. 2020; Powers et al. 2022; Onuț-Brännström et al. 2023; Záhonová et al. 2023). In the case of conflicting results, published information was prioritized over our own analyses.

### Phylogenetics

Maximum likelihood phylogenies were reconstructed for HydA, PFO, ACL, and ACS. Homologs from a balanced selection of taxa were identified by querying a local copy of the NCBI nr database (downloaded Nov 7, 2022) with DIAMOND blastp (bacterial and archaeal) or EukProt (eukaryotic; v3; Richter et al. 2022). Sequences were aligned with MAFFT (v7.526; Katoh et al. 2002), and highly variable sites were masked with BMGE (v1.12; Criscuolo & Gribaldo 2010) using BLOSUM thresholds of 30 (PFO) or 35 (HydA, ACL, ACS).

Maximum likelihood phylogenies were reconstructed using IQ-TREE 2 (v2.3.6; Minh et al., 2020), using ModelFinder to determine the best-fitting model of sequence evolution according to Bayesian information criterion (BIC) (Kalyaanamoorthy et al. 2017) amongst a subset of models (-mset LG+C20, LG+C10, LG+C60, LG+C30, LG+C40, LG+C50, LG). Statistical support was assessed with an ultrafast bootstrap approximation (UFBoot2; Thi Hoang et al. 2017) and SH-like approximate likelihood ratio test (Guindon et al. 2010), with 1000 replicates for each. Trees were visualized with TreeViewer (v2.2.0; Bianchini & Sánchez-Baracaldo 2024) and uninformative branches were collapsed.

### Fluorescence Imaging

Cells in live culture were incubated overnight in 100 nM MitoTracker Orange CMTMRos (ThermoFisher Scientific), a membrane-potential-dependent mitochondrial dye. Cells were fixed for fluorescence imaging for 20 min in 4% (v/v) electron microscopy-grade paraformaldehyde in 1.5× marPHEM buffer with 9% sucrose (Montanaro et al. 2016) and washed with distilled water. Samples were mounted in VECTASHIELD PLUS Antifade Mounting Medium with DAPI (4′,6-diamidino-2-phenylindole; Vector Laboratories) and stored in the dark until imaging. Fluorescence imaging was performed using a Nikon C2+ laser-scanning confocal microscope with 405-nm (DAPI) and 561-nm (MitoTracker Orange) excitation at 100× magnification. Differential interference contrast (DIC) imaging was performed using a Nikon Ti-2A inverted microscope at 100× magnification.

## Acknowledgements

This work was supported by a New Frontiers in Research Fund – Exploration grant (NFRFE-2020-01554) awarded to RMRG and CWS, a grant from the Natural Sciences and Engineering Research Council of Canada (Discovery Grant Program; RGPIN-2019-04336) awarded to RMRG, and a European Union’s Horizon 2020 research and innovation programme (grant agreement ERC Starting grant 101078476) awarded to CWS.

## Data Availability Statement

Raw sequence data have been deposited in the NCBI SRA under BioProject ID PRJNA1531106. Transcriptome and metagenome assemblies, protein multiple sequence alignments, and single-gene phylogenetic trees have been deposited in Figshare (https://figshare.com/s/e7a440b7a8d964379a3d).

