## Supplementary Figures and Table Legends for "A hydrogen-producing mitochondrion in an anaerobic eukaryotrophic rhizarian"

**Table S1.** Sequencing, assembly, and quality metrics of the PCE SSF transcriptome and genome, and PCE Brev transcriptome. Metrics include sequencing read numbers before and after decontamination, and BUSCO completeness scores. BUSCOs not detected in PCE SSF are listed, with those associated with aerobic mitochondrial function highlighted in yellow.

**Table S2.** Predicted MRO-targeted proteins of PCE SSF, including predicted localization, targeting scores, top BLAST hits, Exonerate intron scores, relative expression levels, and transcript and protein sequences.

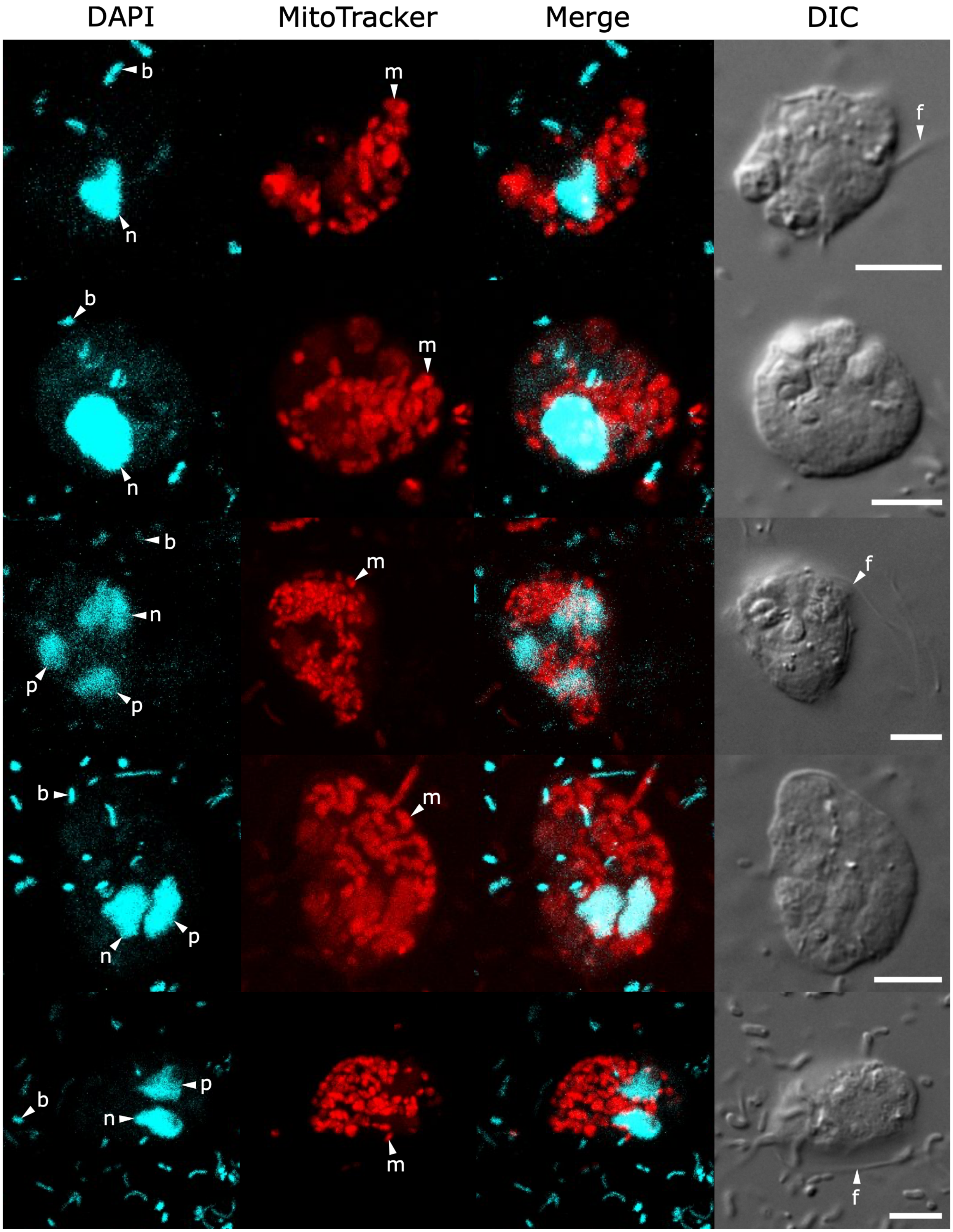


**Fig. S1.** Fluorescence microscopy of additional PCE SSF cells showing DAPI staining of nucleic acids and MitoTracker Orange CMTMRos staining of mitochondrion-related organelles (MROs). Images were acquired with a 100× objective. Scale bars, 5 μm. ‘n’ indicates the nuclear DNA, ‘b’ indicates bacterial DNA, ‘m’ indicates a mitochondrion-related organelle, and ‘f’ indicates a flagellum. In the case of multiple nuclear bodies, the smaller bodies are indicated with ‘p’ to represent putative prey or recently divided nuclei.

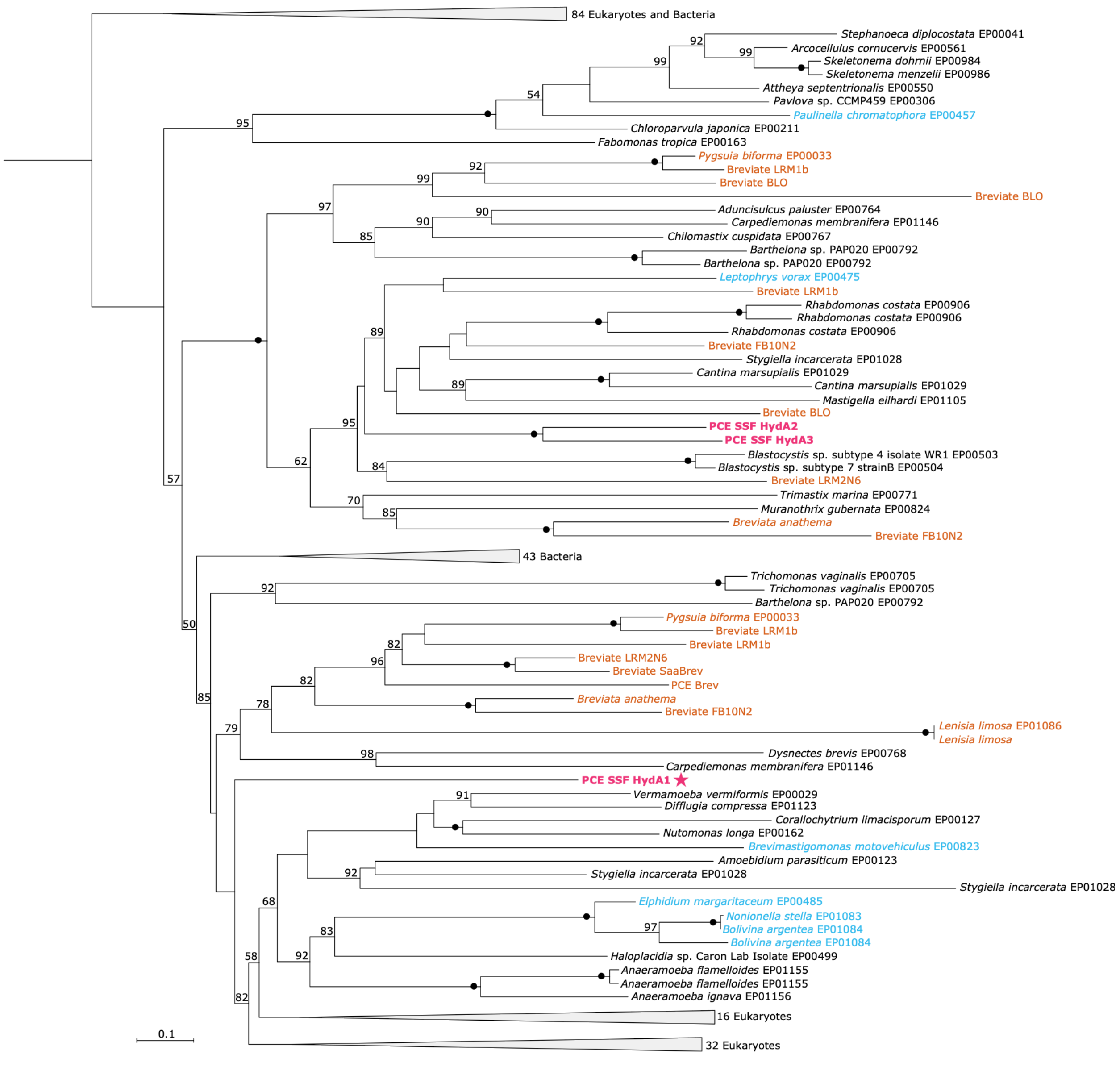


**Fig. S2.** Phylogenetic relationships of PCE SSF [FeFe]-hydrogenases and homologs from other taxa. Multiple alignments were masked using the BLOSUM35 matrix and the phylogeny was inferred with IQ-TREE2 with 245 sequences and 441 sites using the LG+C60+F+R8 model. Branch support was assessed using 1,000 ultrafast bootstrap replicates. Bootstrap values of 100% are indicated with a filled circle, and values below 50% are not shown. Magenta text indicates PCE SSF, blue indicates other rhizarians, and orange indicates breviates. PCE SSF sequences that possess a mitochondrial targeting sequence are indicated by a star.

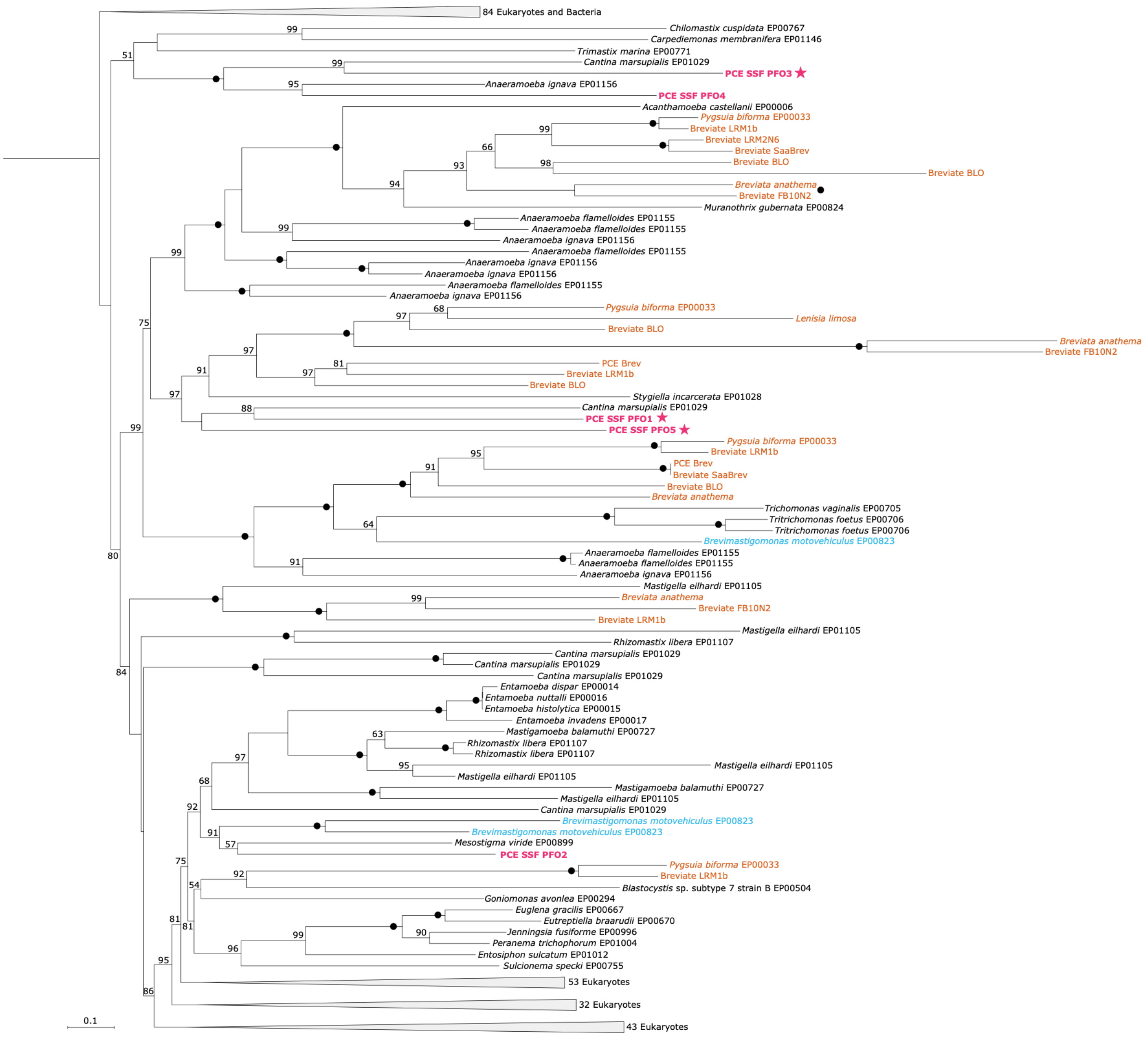


**Fig. S3.** Phylogenetic relationships of PCE SSF pyruvate:ferredoxin oxidoreductase and homologs from other taxa. Multiple alignments were masked using the BLOSUM30 matrix and the phylogeny was inferred with IQ-TREE2 with 407 sequences and 964 sites using the LG+C60+F+I+R10 model. Branch support was assessed using 1,000 ultrafast bootstrap replicates. Bootstrap values of 100% are indicated with a filled circle, and values below 50 % are not shown. Magenta text indicates PCE SSF, blue indicates other rhizarians, and orange indicates breviates. PCE SSF sequences that possess a mitochondrial targeting sequence are indicated by a star.

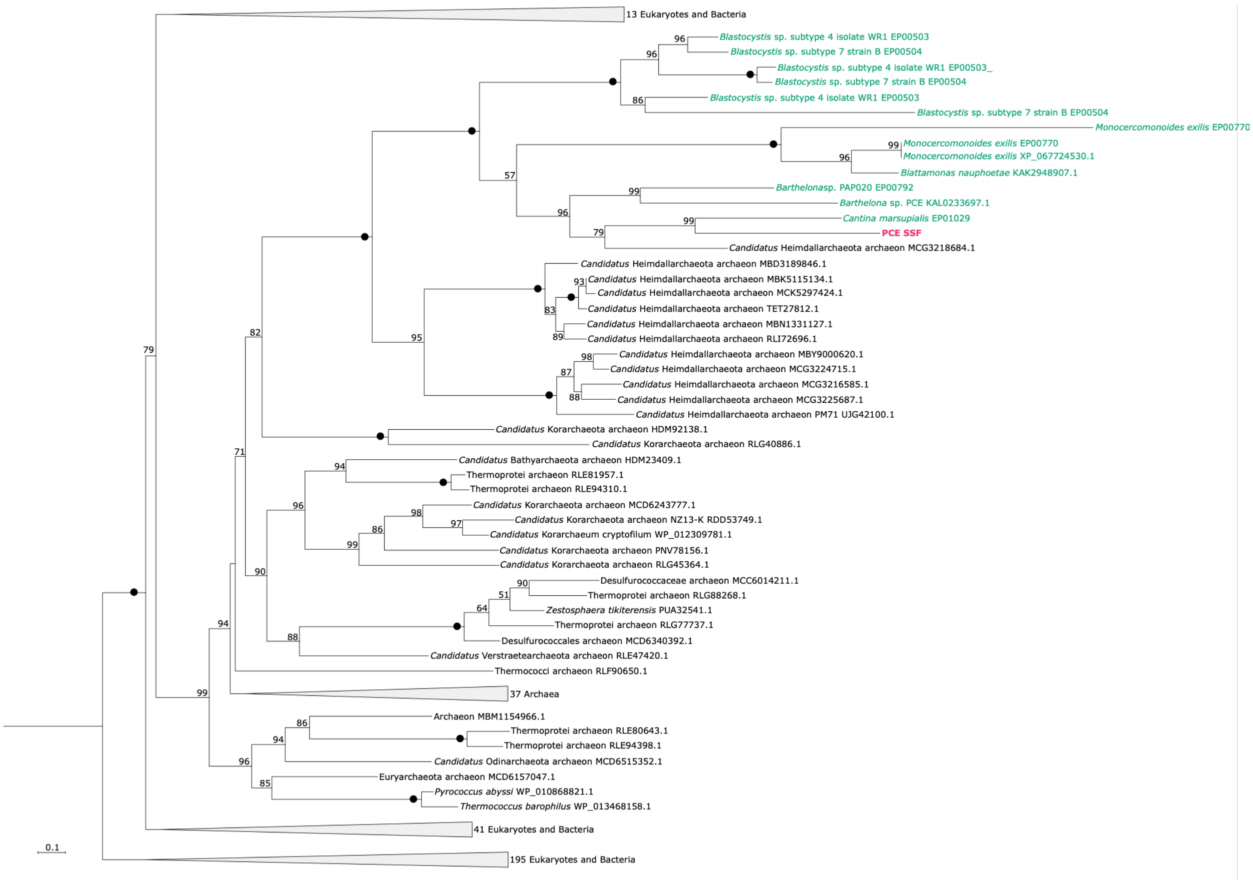


**Fig. S4.** Phylogenetic relationships of PCE SSF acetyl-CoA synthetase and homologs from other taxa. Multiple alignments were masked using the BLOSUM35 matrix and the phylogeny was inferred with IQ-TREE2 with 321 sequences and 274 sites with the LG+C50+F+R9 model. Branch support was assessed using 1,000 ultrafast bootstrap replicates. Bootstrap values of 100% are indicated with a filled circle, and values below 50 % are not shown. Magenta text indicates PCE SSF and teal indicates other eukaryotes.

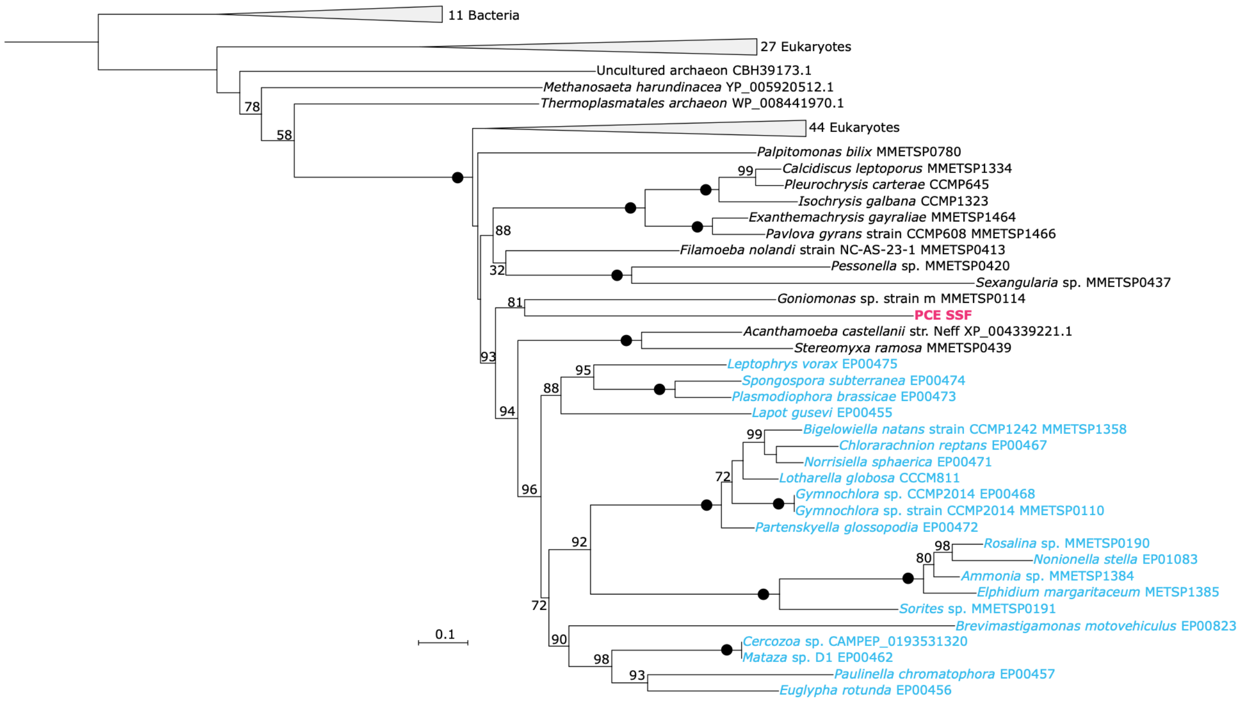


**Fig. S5.** Phylogenetic relationships of PCE SSF ATP-citrate lyase and homologs from other taxa. Multiple alignments were masked using the BLOSUM35 matrix and the phylogeny was inferred with IQ-TREE2 with 125 sequences and 833 sites using the LG+C60+I+R8 model. Branch support was assessed using 1,000 ultrafast bootstrap replicates. Bootstrap values of 100% are indicated with a filled circle, and values below 50 % are not shown. Magenta text indicates PCE SSF and blue indicates other rhizarians.
